# In vitro pancreatic islet cluster expansion facilitated by hormones and chemicals

**DOI:** 10.1101/2019.12.16.873596

**Authors:** Jing-Yu Lin, Jie Cheng, Ya-Qin Du, Wei Pan, Zhong Zhang, Jin Wang, Jie An, Fan Yang, Yun-Fei Xu, Hui Lin, Wen-Tao An, Jia Wang, Zhao Yang, Ren-Jie Chai, Xue-Ying Sha, Hui-Li Hu, Jin-Peng Sun, Xiao Yu

## Abstract

Tissue regeneration, such as pancreatic islet tissue propagation in vitro, could serve as a promising strategy for diabetes therapy and personalized drug testing. However, such a protocol has not been realized yet. Propagation could be divided by two steps, which are: (1) expansion in vitro and (2) repeat passaging. Even the in vitro expansion of the islet has not been achieved to date. Here, we describe a method to enable the expansion of islet clusters isolated from pregnant mice or wild-type rats by employing a combination of specific regeneration factors and chemical compounds in vitro. The expanded islet clusters expressed insulin, glucagon and somatostatin, which are markers corresponding to pancreatic β cells, α cells and δ cells, respectively. These different types of cells grouped together, were spatially organized and functioned similarly to primary islets. Further mechanistic analysis revealed that forskolin in our recipe contributed to renewal and regeneration, whereas exendin4 was essential for preserving islet cell identity. Our results provide a novel method for the in vitro expansion of islet clusters, which is an important step forward in developing future protocols and medium used for islet tissue propagation in vitro. Such method is important for future regenerative diabetes therapies and personalized medicine using large amounts of pancreatic islets derived from the same person.

## Introduction

Diabetes is characterized by insulin resistance and dysfunctional insulin secretion. Recent progress in the generation of functional insulin-producing pancreatic β cells from stem cells differentiation or transdifferentiation in vitro has shed light on cell-replacement therapies for the treatment of diabetes^1–4^. However, compared to tissue/organ replacement therapy derived from somatic islets of the same person for curing diabetes^5,6^, these methods still have the following deficits: (1) The function of a tissue, such as pancreatic islets, cannot be fully substituted by a single β cell type. Islets contain at least three main cell types, and together, they contribute the functional integrity adaptations to responses to different physiological changes^7–10^. In fact, recent reports indicate the essential roles of the islet circuit in maintaining glucose metabolism homeostasis^7,10^. (2) The identity of the β cells generated from transdifferentiation or stem cells is not defined, and their functions are not guaranteed. These cells normally have different gene expression patterns or certain mutations compared to somatic β cells. (3) Some of the β-like cells generated from stem cells may still have pluripotency potential, which may correlate to a risk of tumour development. (4) Finally, these cells may elicit unwanted immune responses.

Ideally, propagating islets isolated from the same patient to the desired number and then transplanting them into the patient is a promising regenerative strategy for diabetes therapy with benefits that stem cell or transdifferentiation based cell-replacement treatment cannot match. This method will minimize the immune response, decrease the identity difference and result in a better ability to adapt to different growth niche after transplantation. In addition, the organoids generated from pancreatic islets propagated from patients can be used for drug testing because they mostly modelling composition, architecture and function of primary tissue. Recently, significant progress has been made in which somatic intestinal, gastric, colonal, hepatic and pancreatic ducts were cultured and propagated to organoids in vitro^11–19^, which lays the foundation for future regenerative therapy. However, these organoids mostly exhibit epithelial properties and are difficult to be induced into endocrine cells. Thus, the in vitro propagation of functional endocrine organoids especially islets, is still required to be established.

Propagation of islets could be divided by two processes: 1) expansion of the islet cells in vitro. 2) Propagating the expanded islet cells by multiple passages. Here, we reported a method for 3D culture with a novel special medium recipe that enabled the in vitro expansion of pancreatic islet clusters isolated from pregnant mice or wild-type rats. Further mechanistic analysis suggested that forskolin (FSK) in the recipe is required for maintaining cell renewal and regeneration, whereas exendin4 is essential for preserving islet identity. Therefore, our results identified a method allowed an in vitro expansion of the pancreatic islet clusters, which could be a key step for propagating patient-specific pancreatic islets for future regenerative diabetes therapy.

## Results

### Recipe for the in vitro pancreatic islet expansion medium (PIEM)

To identify the optimal conditions for pancreatic islet expansion in vitro, we reviewed the chemicals, proliferation and regeneration factors used in the culture of other organoids, such as those in pancreatic ducts, cholangiocytes and hepatocytes^14,16,20,21^, and pancreatic islet organoids derived from stem cells or fibroblasts (Fig. 1)^1–4^. According to their reported functions, these chemicals and factors are classified into the following groups: islet identity, cell division and proliferation, and cell renewal and regeneration (Fig. 1). By screening different combinations of compounds and factors, a medium composed of 12 chemicals and hormones, as well as the general nutrient nicotinamide, B27 and GlutaMAX, was created and was found to robustly support the in vitro expansion of dispersed islet clusters isolated from pregnant mice or wild-type rats. We named this in vitro pancreatic islet expansion medium PIEM.

**Fig. 1.**
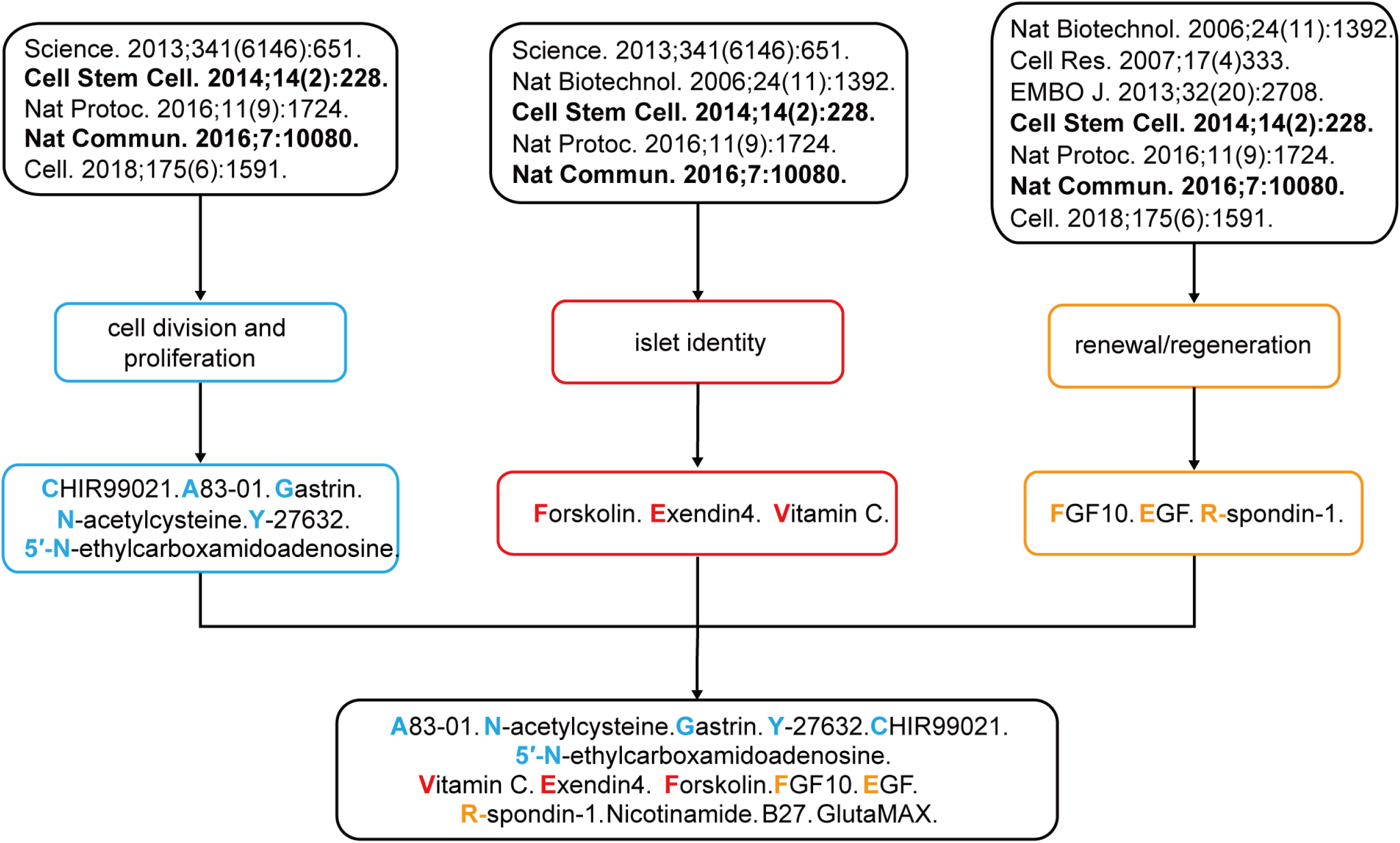
Flowchart for the development of the medium recipe used for pancreatic islet expansion. The chemicals and hormones used in previous studies for stem cell expansion, differentiation and pancreatic islet cell conversion (upper panel) are classified by their reported function related to cell division and proliferation, islet identity, and cell renewal and regeneration (middle panel). The common factors are summarized and combined for the formulation of the medium recipe used for in vitro pancreatic islet cluster expansion.

### Expansion of mouse islet clusters in vitro

Previous studies have shown that pregnant mouse islets maintain expansion ability in vivo, whereas wild-type mouse islets undergo little expansion^22,23^. We therefore used islets from both pregnant and wild-type mice, as pregnant mice could serve as an easier starting point for investigating the regeneration process. We isolated pancreatic islets from both wild-type and pregnant C57BL/6 mice by collagenase P perfusion, sedimentation and handpicking. These pancreatic islets were separated into single cells or cell clusters by dispase II digestion and mechanical dissociation. The single cells or cell clusters were cultured in 3-dimensional (3D) Matrigel. Only dispersed islet clusters isolated from pregnant mice showed significant expansion during nine-day culture (Fig. 2a). Approximately 5~15% of the pancreatic islet clusters isolated from pregnant mice were able to expand from a surface area of 6,000-9,000 μm^2^ (240-320 cells) to a surface area of 20,000-30,000 μm^2^ (1300-2100 cells) (Fig. 2a, c). In contrast, the dispersed single islet cells from pregnant or wild-type mice or islet clusters from wild-type mice showed no significant expansion (Fig. 2b, d).

**Fig. 2.**
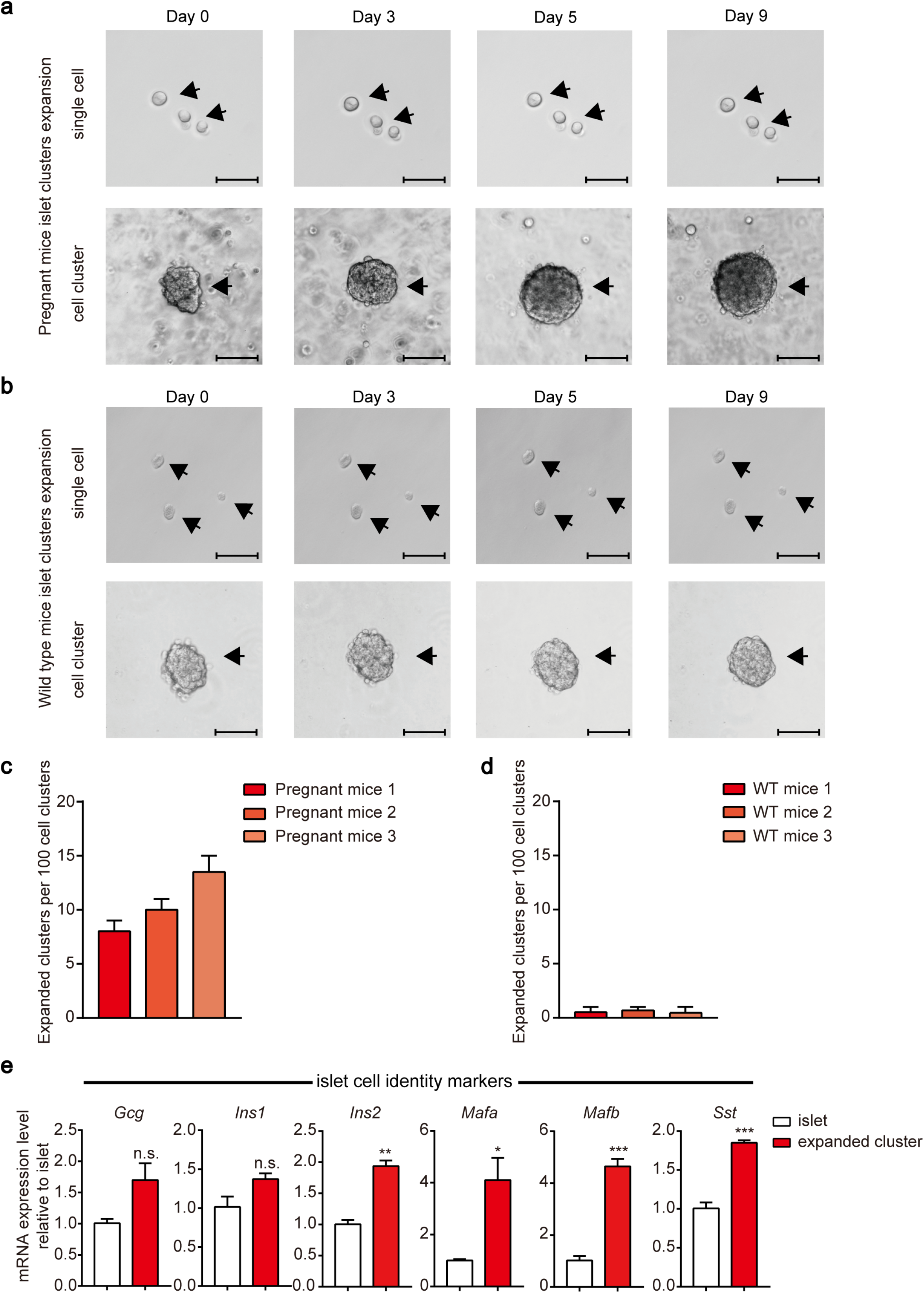
In vitro expansion of pregnant mouse islet clusters. **a** Representative images of single islet cells or islet clusters (approximately 6,000-9,000 μm^2^ original surface area) derived from pregnant mice after 3D culture in pancreatic islet cluster expansion medium (PIEM) at the indicated time points. Experiments were performed on pregnant mice. The scale bar in the single-cell image represents 50 μm, and one in the cluster image represents 100 μm. **b** Representative images of dispersed wild-type mouse islet single cells or clusters at various times after 3D culture with PIEM. The scale bar in the single-cell image represents 50 μm, and one in the cluster image represents 100 μm. **c** Numbers of expanded clusters per 100 islet cell clusters. The experiments were performed in triplicate and with pregnant mice. The data are represented as the mean ± SEM. **d** Numbers of expanded clusters per 100 islet cell clusters. The experiments were performed in triplicate on wild-type mice. The data are represented as the mean ± SEM. **e** Comparison of islet cell identity marker expression in pregnant mouse pancreatic islet clusters vs. isolated primary mouse islets. *, p <0.05; **, p < 0.01; ***, p < 0.001; clusters were compared with primary mouse islets. The data are shown as the mean ± SEM of at least three independent experiments. The data statistics were analysed using an unpaired two-tailed Student’s t-test.

We next used qRT-PCR to examine the expression of key islet markers in the expanded islet clusters. The expanded islet clusters maintain expression of the crucial pancreatic β cell markers *insulin-1 (Ins1), insulin-2 (Ins2)* and *Mafa*; the α cell markers *glucagon (Gcg)* and *Mafb*; and the δ cell marker *somatostatin (Sst)* (Fig. 2e).

To confirm that specific cell types coexisted in the in vitro expanded islet clusters, which is a hallmark of expanded islet clusters, we performed immunostaining for INSULIN, GLUCAGON and SOMATOSTATIN, which are markers for pancreatic α cells, β cells and δ cells, respectively. The result unambiguously identified that glucagon, insulin, and somatostatin staining was present in different cells grouped together in the same expanded islet clusters similar to primary islets (Fig. 3a, b and Supplementary Fig. S1a, b). The percentage of the α cells, β cells and δ cells are (12.4±3.67)%, (78±4.55)%, (7.68±1.26)% respectively in propagated islet clusters, which is similar to the pancreatic islets isolated from pregnant mice (Fig. 3c). Importantly, we also observed the co-localization of the pancreatic transcription factors PDX1 and MAFA with the insulin, as well as the partial co-localization of the NKX6.1 with the insulin in the same cells of expanded islet clusters, which is similar to the pancreatic islets isolated from pregnant mice(Fig. 3d-f and Supplementary Fig. S1c-e).

**Fig. 3.**
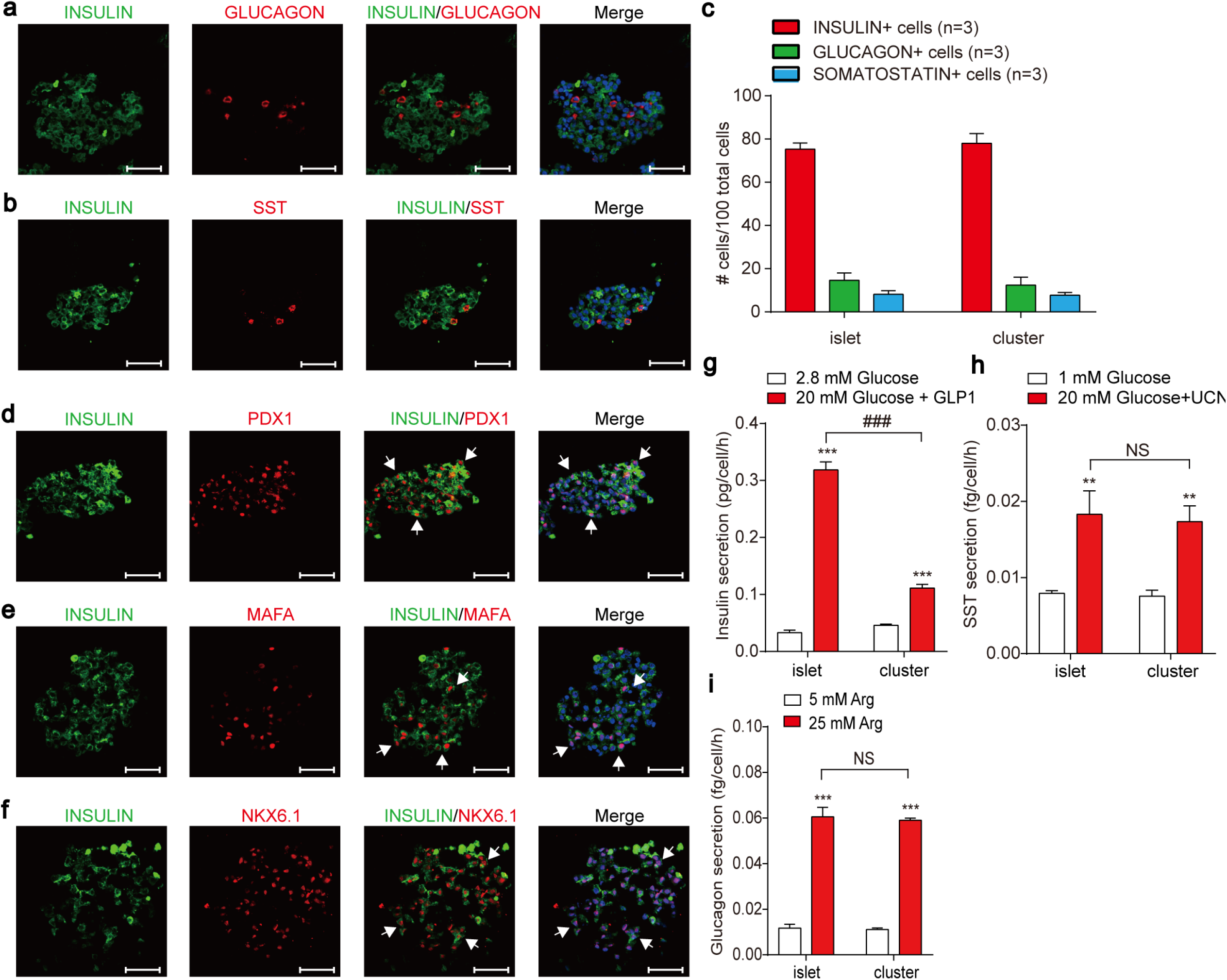
Immunostaining for pancreatic islet cell markers and functional examination of in vitro expanded pancreatic islet clusters. **a** Immunofluorescence staining for INSULIN and GLUCAGON in expanded pancreatic islet clusters derived from pregnant mice; the scale bar represents 50 μm. **b** Immunofluorescence staining for INSULIN and SOMATOSTATIN in expanded pancreatic islet clusters derived from pregnant mice; the scale bar represents 50 μm. **c** Cell composition of expanded pancreatic islet clusters. Error bars represent mean ±SD (n=3) **d** Immunofluorescence staining for INSULIN and PDX1 in expanded pancreatic islet clusters derived from pregnant mice; the scale bar represents 50 μm. **e** Immunofluorescence staining for INSULIN and MAFA in expanded pancreatic islet clusters derived from pregnant mice; the scale bar represents 50 μm. **f** Immunofluorescence staining for INSULIN and NKX6.1 in expanded pancreatic islet clusters derived from pregnant mice; the scale bar represents 50 μm. **g** Insulin secretion of primary islets or expanded islet clusters treated with 20 mM glucose and 100 nM glucagon peptide 1 (GLP-1) were measured for 30 minutes. The control groups were treated with 2.8 mM glucose. ***, p < 0.001, stimulation groups were compared with control groups. ###, p < 0.001, expanded islet clusters were compared with primary islets. **h** Somatostatin secretion of primary islets and expanded islet clusters treated with 20 mM glucose and 100 nM UCN3 were measured for 1 hour. The control groups were treated with 1 mM glucose. **, p < 0.01, stimulation groups were compared with control groups. NS, no significance, expanded islet clusters were compared with primary islets. **i** Glucagon secretion of primary islets and expanded islet clusters treated with 25 mM Arginine were measured for 1 hour. The control groups were treated with 5mM Arginine. ***, p < 0.001, stimulation groups were compared with control groups. NS, no significance, expanded islet clusters were compared with primary islets. The data are shown as the mean ± SEM of at least three independent experiments. The data statistics were analysed using an unpaired two-tailed Student’s t-test.

We next examined the functional integrity of the expanded islet clusters. In response to combined stimulation with 20 mM glucose and 100 nM GLP-1, the expanded islet clusters demonstrated a significant increase in insulin secretion, although with a smaller extent compared to the pregnant mouse islets (Fig. 3g). In response to high glucose stimulation, both other labs and our lab have shown that UCN3 serves an endogenous paracrine factor secreted by pancreatic β cells to stimulate endogenous somatostatin secretion from pancreatic δ cells^7,10,24^. We therefore stimulated the expanded islet clusters with both glucose and UCN3 to minimize the amount required for islet cluster usage. The in vitro expanded islet clusters displayed significantly more somatostatin release in response to glucose and UCN3 stimulation (Fig. 3h). Moreover, the expanded islet clusters secreted more glucagon in response to 25mM Arginine stimulation, hallmarking the normal function of pancreatic islet α cells, which is similar to the pregnant mouse islets (Fig. 3i). These results confirmed the functional integrity of the in vitro expanded islet clusters isolated from pregnant mice.

Although the expanded pancreatic islet clusters showed similar SST and Glucagon secretion compared to the primary isolated islets in response to specific physiological stimulations, it is worth to note that their insulin secretion in response to the combined stimulation of glucose and GLP-1 is significantly weakened. Some extent degeneration during the islet expansion may occur during current in vitro expansion condition, which awaits for further investigation and methods optimization in our future work.

### Expansion of rat islet clusters in vitro

We then examined the expansion of the dispersed rat islet single cells and clusters in PIEM. Similar to those from pregnant mice, approximately 5~15% of the rat pancreatic islet clusters were able to expand from a surface area of 5,000-8,000 μm^2^ (200~350 cells) to a surface area of 20,000-25,000 μm^2^ (1,000~1,900 cells), whereas dispersed rat islet single cells showed no such expansion ability (Fig. 4a, b). Furthermore, qRT-PCR revealed that the expanded rat islet clusters maintained the same expression of the α cell markers *Gcg* and *Mafb* as the primary rat islets but showed significantly higher expression of the β cell markers *Ins-1, Ins-2* and *Mafa* and the δ cell marker *Sst* (Fig. 4c).

**Fig. 4.**
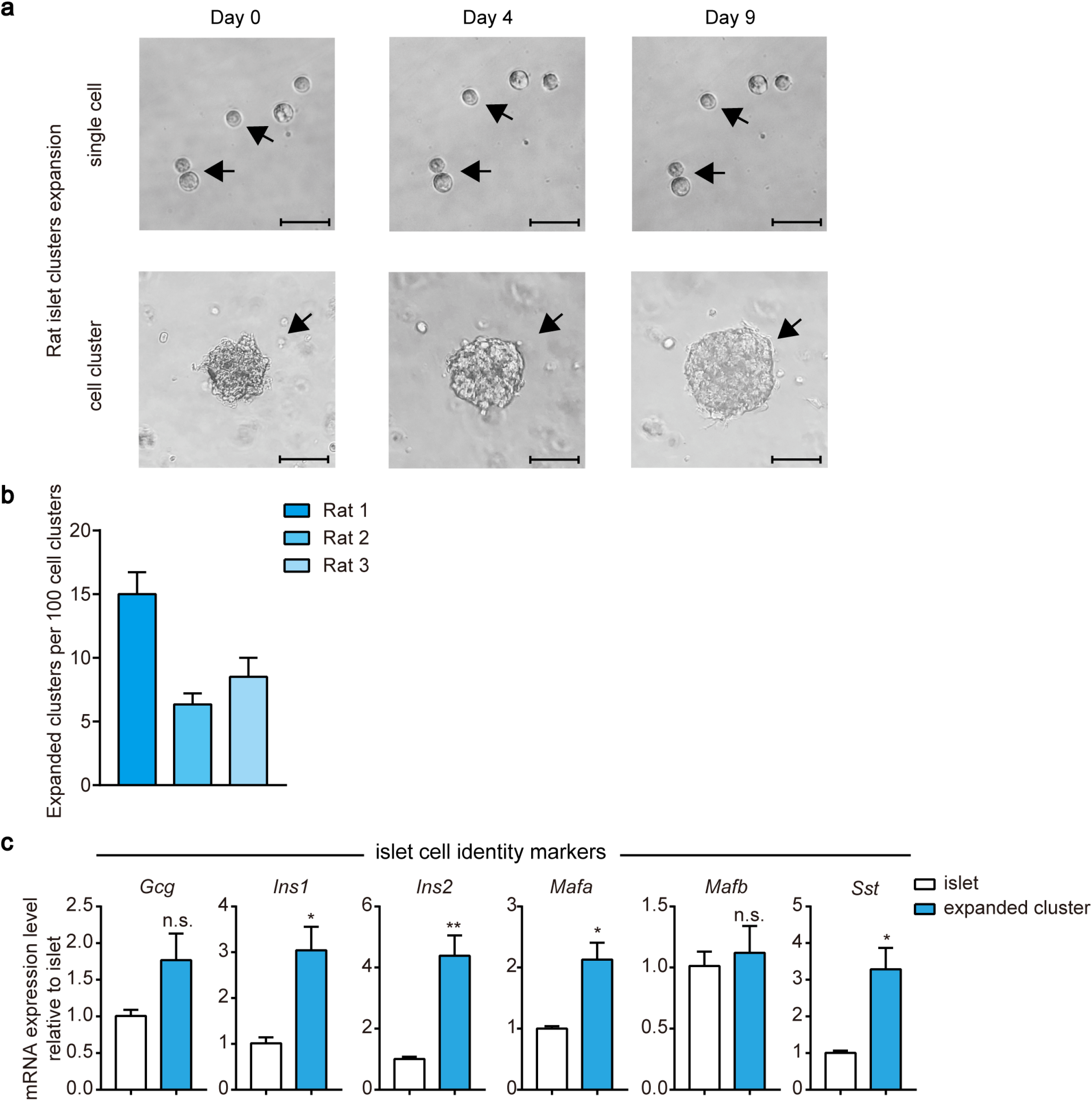
In vitro expansion of rat islet clusters. **a** Representative images of rat islet single cells or clusters (approximately 5,000-8,000 μm^2^ original surface area) at various times after 3D culture with PIEM. The scale bar in the single-cell image represents 50 μm, and one in the cluster image represents 82.5 μm. **b** Numbers of expanded clusters per 100 islet cell clusters. The experiments were performed on rats in triplicate. The data are represented as the mean ± SEM. **c** Comparison of the expression of islet cell identity markers in rat expanded clusters compared with primary islets. *, p <0.05; **, p < 0.01; clusters were compared with primary rat islets. The data are shown as the mean ± SEM of at least three independent experiments. The data statistics were analysed using an unpaired two-tailed Student’s t-test.

### Increased gene expression related to dedifferentiation, pluripotency and proliferation in expanded islet clusters

We next examined whether the expanded islet clusters increased the expression level of genes functionally associated with cell proliferation, renewal and regeneration, which are important factors for in vitro regeneration. Importantly, significantly higher expression of *Ki67*, *Ccnd1* and *Pcna* was found in expanded islet clusters than in islets derived from pregnant mice or wild-type rats, highlighting the increased proliferation ability of the expanded islet clusters (Fig. 5a, d). Moreover, significantly higher *Nanog* and *Sox9* expression were found in expanded islet clusters derived from both pregnant mice and rats (Fig. 5b, e). These results provide putative explanations accounting for the better proliferation and renewal abilities of isolated islet clusters than integral islets.

**Fig. 5.**
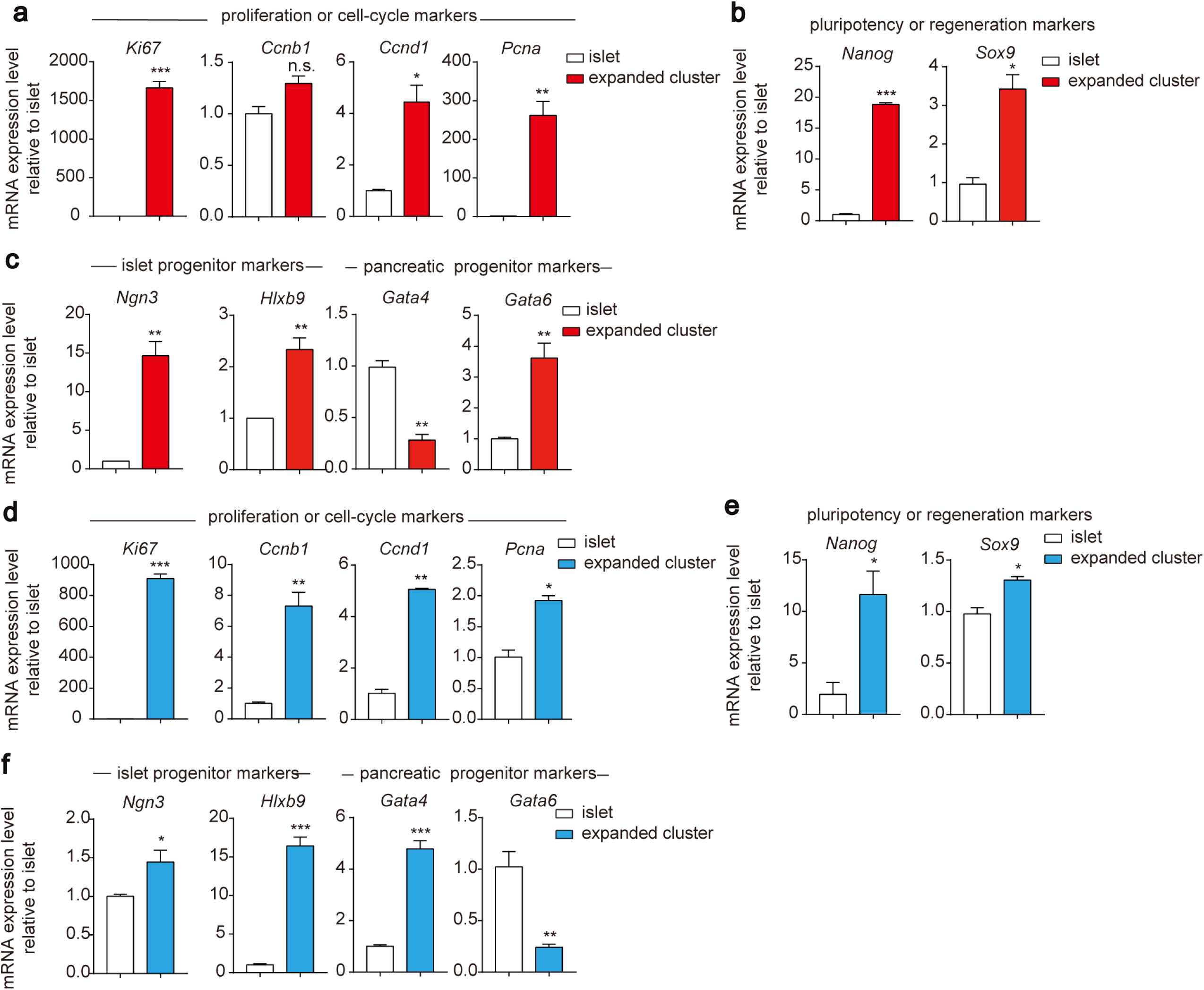
qRT-PCR analysis of gene expression in expanded clusters compared to primary islets. **a** Comparison of the expression of proliferation and cell cycle markers in pregnant mouse expanded clusters vs. isolated primary islets. **b** Comparison of the expression of pluripotency and regeneration markers in pregnant mouse expanded clusters vs. isolated primary islets. **c** Comparison of the expression of islet and pancreatic progenitor markers in pregnant mouse expanded clusters vs. isolated primary islets. **d** Comparison of the expression of proliferation and cell cycle markers in rat expanded clusters vs. isolated primary islets. **e** Comparison of the expression of pluripotency and regeneration markers in rat expanded clusters vs. isolated primary islets. **f** Comparison of the expression of islet and pancreatic progenitor markers in rat expanded clusters vs. isolated primary islets. *, p < 0.05; **, p < 0.01; ***, p < 0.001; clusters were compared with primary islets. The data are shown as the mean ± SEM of at least three independent experiments. The data statistics were analysed using an unpaired two-tailed Student’s t-test.

We suspected that these islets gained proliferation and pluripotency due to dedifferentiation. Thus, we examined dedifferentiation markers in expanded islet clusters and compared them with those in isolated pancreatic islets. Interestingly, the expanded islet clusters showed significantly higher expression of *Ngn3* and *Hlxb9*, markers that characterize pancreatic islet progenitor cells (Fig. 5c, f). Although the expanded islet clusters derived from pregnant mice showed higher expression of *Gata6*, they showed decreased expression of *Gata4*, both of which are pancreatic progenitor markers (Fig. 5c, f). The protein expression of KI67 and SOX9 in expanded islet clusters were confirmed by immunofluorescence, higher than in primary islets (Supplementary Fig. S2a-d). Similarly, the expanded islet clusters derived from rats did not display significant pancreatic progenitor characteristics, as they showed decreased expression of *Gata6* (Fig. 5c, f). The data suggested that these expanded islet clusters underwent one-step dedifferentiation towards islet progenitor cells, but they did not ultimately reach the pancreatic progenitor cell stage. Taken together, this one-step dedifferentiation of isolated islet clusters cultured in PIEM towards pancreatic islet progenitor cells contributed to the gain of cell proliferation, renewal and regeneration functions of expanded islet clusters.

### Essential role of FSK and exendin4 in PIEM

During our formula component screening and recipe formation for PIEM, we identified that FSK and exendin4 are both required for robust islet cluster expansion in vitro. It is known that exendin4 induced cAMP accumulation after it activates GLP-1R and downstream Gs proteins^25^. FSK is known as a cAMP agonist through direct binding to adenyl cyclase^26^. Moreover, recent reports indicate that FSK is required for proliferation of liver organoids in addition to A83-01^27^. Interestingly, removing FSK or exendin4 from PIEM has different effects on the gene expression profiles of islet identity, proliferation, renewal and regeneration markers. Removing FSK from PIEM had partial effects on islet identity and cell proliferation markers, including decreased expression of *Mafa*, *Sst* and *Ccnb1* (Fig. 6a, b). In particular, FSK was required for the expression of cell renewal and regeneration markers, including *Nanog* and *Sox17* (Fig. 6c). In contrast, exendin4 was essential for islet identity and proliferation because it maintained the expression of the pancreatic β cell markers *Ins-1*, *Pdx1* and *Mafa* and the proliferation markers *Ki67*, *Ccnb1* and *Cdk4* (Fig. 6a, b). In particular, exendin4 was required for the expression of *Ngn3* and *Hxlb9*, two islet progenitor markers (Fig. 6d). These results indicated that FSK in PIEM renders cell renewal and regeneration ability to isolated islet clusters, whereas exendin4 is essential for maintaining islet β cell identity, proliferation and dedifferentiation.

**Fig. 6.**
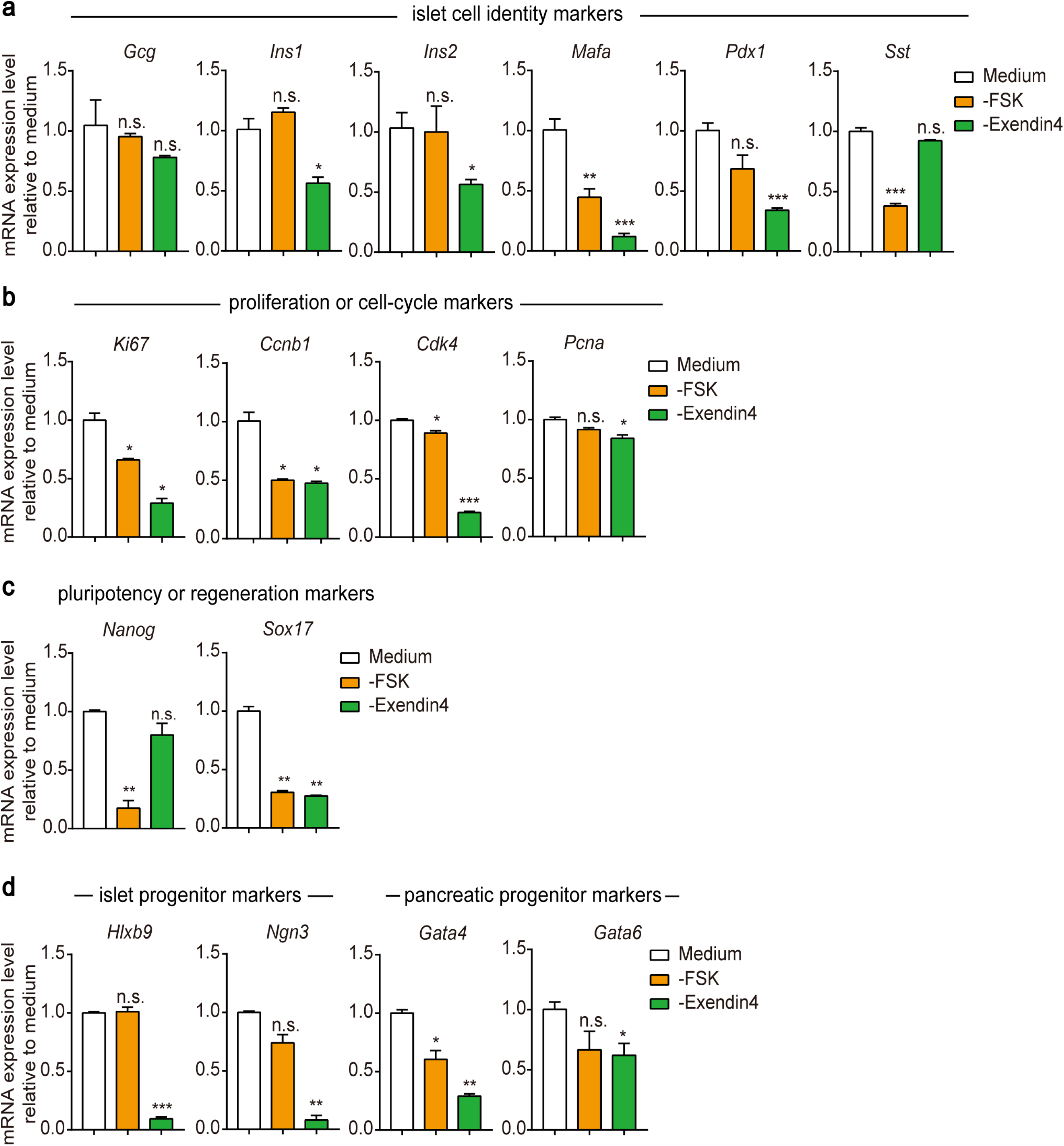
Essential roles of FSK and exendin4 in in vitro pancreatic islet expansion. **a-d** qRT-PCR analysis of the expression of islet cell identity markers (**a**), proliferation and cell cycle markers (**b**), pluripotency and regeneration markers (**c**) and islet and pancreatic progenitor markers (**d**) in pregnant mouse clusters after 3 days of culture with PIEM or PIEM without forskolin or exendin4. *, p < 0.05; **, p < 0.01; ***, p < 0.001; cell clusters cultured in PIEM without FSK or exendin4 were compared with those cultured in complete PIEM. The data are shown as the mean ± SEM of at least three independent experiments. The data statistics were analysed using an unpaired two-tailed Student’s t-test.

## Discussion

The in vitro expansion of primary pancreatic β cells or pancreatic islet tissues has not been reported before. Here, we developed a medium recipe, which we named PIEM, that enabled the proliferation of dispersed islet clusters isolated from pregnant mice or wild-type rats in vitro (Fig. 7). Isolated islet clusters normally grow from an initial surface area of 5,000-9,000 μm^2^ to a surface area of 20,000-30,000 μm^2^, with an estimated 5~8-fold increase in cell number. Compared to primary pancreatic islets, expanded cell clusters have similar or higher mRNA expression levels of insulin, glucagon and somatostatin, which are markers of pancreatic β cells, α cells and δ cells, respectively. Expanded islet clusters demonstrated normal insulin and somatostatin secretion in response to physiological stimulation and normal glucagon secretion in response to amino acids, suggesting that the clusters recapitulate specific functions of pancreatic islets. Immunostaining further confirmed that the cells expressing insulin, glucagon and somatostatin were grouped together and spatially organized in expanded islet clusters similar to primary islets. These data indicated that the expanded islet clusters behave similar to an in vitro organoid, thus serving as an initial step for future islet propagation methods developing in vitro. The ultimate goal of developing efficient methods for propagation islets in vitro is to fulfil the demand for regenerative diabetes therapy and personalized drug tests.

**Fig. 7.**
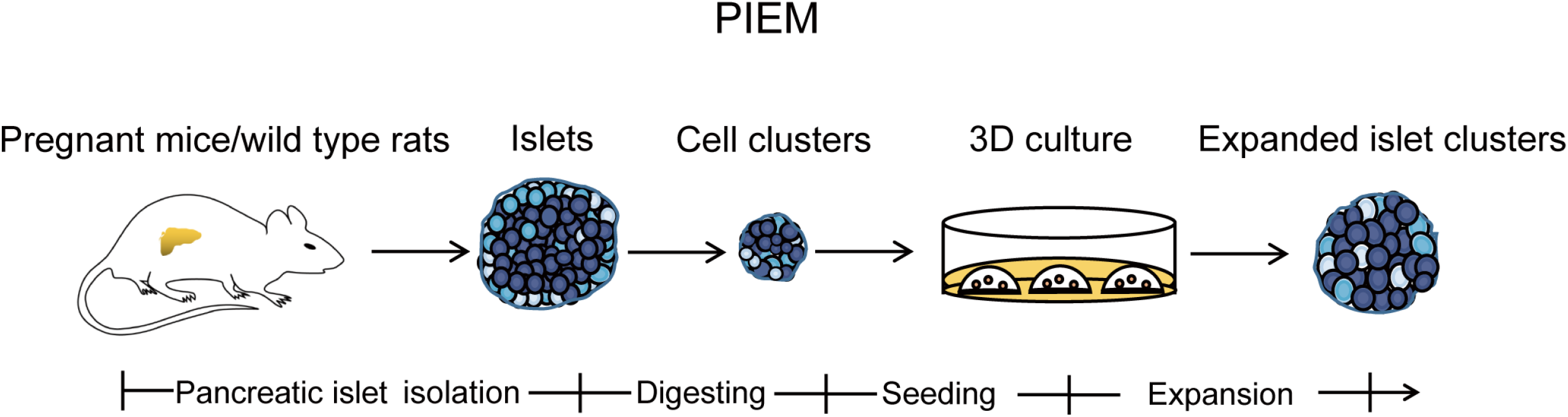
Scheme of the current approach and future direction. Schematic depicting the isolation, seeding and the expansion of primary islet cell clusters. After the islets were isolated from pregnant mice or wild-type rats, the islets were digested into appropriate cell clusters by incubation with the dispase II. The digestion time were optimized and the mechanical blow force was used for clusters generation. These clusters were cultured 3D in PIEM to achieve the in vitro expansion of the islet cell clusters.

It is worth noting that we observed the expansion of islet clusters isolated from only pregnant mice and wild-type rats in PIEM, not those isolated from wild-type mice. Pregnant mice are well known for their increased pancreatic β cell proliferation^28–31^, which provides a useful starting point for testing medium recipes. Although we did not observe the proliferation of islet clusters isolated from wild-type mice, we did observe the expansion of islet clusters isolated from wild-type rats, which share more properties identical to human islets than mouse islets. An urgent need is to test whether PIEM enables the expansion of primary islet clusters isolated from human patients.

Another notable observation is that only dispersed islet clusters were able to proliferate in vitro, whereas dispersed single primary islet cells could not. Two important factors may contribute to this discrepancy. First, the islet clusters contain multiple cell types, which are not only the origins of the different cell types in the finally expanded islet clusters but also may form certain cell circuits and gradient hormone concentrations to support in vitro expansion. Second, primary single cells must undergo harsher digestion than islet clusters, which means that the extracellular parts of the membrane or matrix proteins, including receptors, ion channels and transporters, may be digested and their function impaired. These membrane or matrix proteins may be key for in vitro tissue expansion. Therefore, future analysis of the different expression patterns and functions of membrane proteins and signalling circuit differences between isolated islet clusters and single cells may provide more clues for unravelling the secrets of the in vitro expansion of islets and provide guidance for a better strategy for in vitro islet cluster expansion.

We noticed that expanded islet clusters have higher expression of *Ngn3* and *Hlxb9*, which are markers of pancreatic islet progenitor cells. Although the *Nanog* in islet clusters increased approximately 10 fold compared to normal islets, however, this level is more mimic the whole embryo tissue but much lower (more than 1,000 fold lower) than the embryo stem cells (Supplementary Fig. S2e). Whether this amount *Nanog* is required for islet expansion and propagation in vitro, and whether it has cancer potential require further investigation. This observation indicated that in vitro expanded islets underwent certain dedifferentiation. However, we doubt that the islet clusters gained proliferation ability through this dedifferentiation. Further experiments with withdrawal of key chemicals in the medium will test this hypothesis. For clinical usage, the in vitro expansion of 100 patient islets to 1 million islets without significant gene mutations, while preserving pancreatic islet function, and transferring them back to the patient to recover glucose homeostasis is ideal. However, the current approach of PIEM allows only 5~8-fold growth of approximately 5%-15% of isolated islet clusters. The expansion of the islet clusters was only observed in the first passage. Therefore, it is unlikely to cause significant genomic instability. However, the current acquired in vitro expanded islet cluster cells do not contain sufficient amounts for RNA-seq to verify the gene expression profile or transplantation to confirm their in vivo functions. More important, the final usage of in vitro propagated islet clusters for regenerative therapy to treat diabetes in general require millions of pancreatic islets. Therefore, a future revision of the current PIEM composition and method for in vitro islet propagation is in an urgent demand.

One of the future directions for the optimization of in vitro islet propagation methods could be as follows: (1) initial expansion and dedifferentiation of islet clusters to islet progenitor-like cell clusters, (2) rapid proliferation of islet progenitor-like cell clusters to a sufficient amount, (3) differentiation of progenitor-like cell clusters to functional integral islets, (4) transplantation or (5) exploitation of these expanded islet tissues for drug tests. Therefore, manifesting the markers for each stage and searching chemicals that are useful for each procedure in the above protocol is our next goal to fulfil expanded islet clusters regeneration in vitro in the future.

Taken together, we have developed a recipe for PIEM, and our 3D culture method enabled the initial generation of expanded islet clusters via the in vitro expansion of isolated islet clusters from pregnant mice or wild-type rats in the first passage. Our study is one important step forward for the realization of a method to fulfil in vitro propagation of pancreatic islet in vitro, which may facilitate new therapies development to treat diabetes.

## Materials and methods

### Animals

C57BL/6 mice and rats of Wistar strain were housed and bred under specific-pathogen-free conditions at Shandong University animals care facility. Female animals were used. The numbers of animals are indicated in the figure legends within each experiment. All animal care and experiments were reviewed and approved by the Animal Use Committee of Shandong University School of Medicine.

### Pancreatic islet isolation and digestion into single cells or cell clusters

We isolated pancreatic islets from C57BL/6 mice, pregnant C57BL/6 mice and Wistar rat. Briefly, mice and rat were killed by cervical dislocation and then pancreas were isolated from them individually. Adult pancreas were digested by Collagenase P (Roche, 11213873001) at 37□ for 18-25 minutes. Digestion was stopped by cold Hank’s balanced salt solution (136.9 mM NaCl; 5.4 mM KCl; 1.3 mM CaCl_2_; 0.8 mM MgSO_4_; 0.44 mM KH_2_PO_4_; 0.34 mM Na_2_HPO_4_; 5.55 mM D-glucose; 4.4 mM NaHCO_3_, pH=7.4) followed by sedimentation for three or five times at 4□. The islets were collected by hand picking using a stereomicroscope and were cultured overnight in islet complete media containing 5.6 mM glucose (Biological Industries), 10% FBS (Gibco),0.1% penicillin/streptomycin.Next we obtained single cells and cell clusters of different diameters (20-150μm)under the microscope by controlling enzyme concentration, digestion and settling time. Islets briefly after naturally settling for 1 minute in KRBB buffer (135 mM NaCl; 4.7 mM KCl; 1.2 mM KH_2_PO_4_; 10 mM Hepes; 3 mM D-glucose; 5 mM NaHCO_3_; 0.1% BSA; 1% penicillin/streptomycin, pH=7.4), preheated EDTA-KRBB solution was added, gently piped for 2 min, then placed in a 33 □ water bath for 10 min; after clearing the supernatant, digested with dispase II(0.1-0.5 U/ml, Roche, 04942078001) in a 33 □ water bath for 3~8 min to obtain single cells or islet cell clusters. Mechanical fragmentation was achieved by gentle pipetting (5–10 times) with a glass dropper in a 15ml glass centrifuge tube. Using DMEM/F-12 (Biological Industries) medium with 10% FBS (Gibco) and 1% penicillin/streptomycinfor termination of digestion.Collecting single cells or cell clusters by centrifugation (300×g, 5 min, Room temperature).

### Production of Rspo1-conditioned medium

The RSPO1 conditioned medium is home-made and will be described in other papers.

### Expanded islet clusters culture

Isolated islet cell clusters were washed twice with DMEM/F-12 (Biological Industries), counted and mixed with Matrigel in glass centrifuge tubes. 30,000–50,000 cells or 10-20 cell clusters were used per well of a pre-warmed 24-well plate (the volume of 50 μl). After Matrigel was solidified for 30min at 37□, pancreatic islet expansion medium (PIEM) was added. PIEM consists of DMEM/F-12 (1% GlutaMax,1% Penicillin-Streptomycin) plus 15% RSPO1 conditioned medium (home-made), 3 μM CHIR-99021 (BioGems), 1 μM A83-01 (Adooq Bioscience), 10 nM gastrin-1 (MedChemExpress), 1.25 mM N-acetylcysteine (Sigma), 10 μM Y-27632 (Adooq Bioscience), 10 μM forskolin (TargetMol), 50 ng/ml Exendin-4 (ChinaPeptides), 50 μg/ml L-Ascorbic acid (Sigma), 250 nM 5-Iodotubercidin (Adooq Bioscience), 50 ng/ml FGF10 (PeproTech), 50 ng/ml EGF (PeproTech), 10 mM Nicotinamide (Sigma), B27 Supplement (minus Vitamin A) (Thermo). Cultures were kept at 37□, 5% CO_2_ in a humidified incubator. During culturing, medium was refreshed every three days.

### RNA extraction and qRT-PCR

RNA from islets and islet single cells as well as islet cell clusters derived from C57BL/6 mice, pregnant C57BL/6 mice and Wistar rat, Including the corresponding islet clusters were extracted with TRIzol reagent (Thermo). We used the tip to pick up the expanded islet clusters from the Matrigel using a stereomicroscope. Cell Recovery Solution(Corning) is used to recover cells from Matrigel. Add 0.2-0.5 ml per 24-well plate cold Cell Recovery Solution. Scrape the gel layer into a cold 1.5 ml Microcentrifuge Tube and leave it on ice for 0.5-1 hour until the gel complete dissolved. Collecting cells by centrifugation (200×g, 5 min, 4□). And we used TURBO DNA-free TMKit (Invitrogen) to remove DNA contamination from the extracted RNA. cDNA synthesis using the qRT-PCR Kit (Toyobo, FSQ-101). We conducted quantitative reverse-transcriptase PCR (qRT-PCR) in the LightCyclerqPCR apparatus (Bio-Rad) with the FastStart SYBR Green Master (Roche). All primers used for qRT-PCR assay are listed in Supplementary Table S1.

### Immunofluorescence staining

Expanded islet clusters were harvested using cell recovery solution (Corning, 354253) and fixed in 4% paraformaldehyde for 30 min at room temperature, followed with blocking and permeabilizing in PBS with 0.5% Triton X-100 (Solarbio, T8200) and 5% donkey serum (Solarbio, SL050) for 30min at room temperature. Then, samples were incubated with primary antibody at 4°C overnight, followed by incubation with secondary antibody for 2h at room temperature. DAPI (Beyotime, C1002) was used to stain the nucleus and find islets. The following antibodies were used for immunofluorescence: anti-INSULIN (1:200, sc-9168; Santa), anti-SOMATOSTATIN (1:600, ab30788; Abcam), anti-GLUCAGON (1:200, G2654; Sigma). anti-PDX1 (1:200, ab47267; abcam), anti-SOX9 (1:200, ab185966; abcam), anti-NKX6.1 (1:200, ab221549; abcam), anti-MAFA (1:200, ab26405; abcam), anti-KI67 (1:200, D3B5; Cell Signaling Technology). Expanded islet clusters imaging was performed on Zeiss LSM 780 and processed using ImageJ or Adobe illustrator software.

### Insulin secretion measurement

The insulin secretions were preformed similar to previously described in our group^7,9,24^. Fifty day 7 expanded islet clusters (cell number is comparable to 1/10 of primarily isolated islets, 1,000~2,000 cells) or ten islets for each group were starved in 2.8 mM glucose MKRBB buffer (5 mM KCl, 120 mM NaCl, 15 mM Hepes, 24 mM NaHCO3, 1 mM MgCl2, 2 mM CaCl2, pH=7.4) for 1 hour at 37°C, and then treated with 20 mM glucose and 100 nM GLP1, whereas the control groups were treated with 2.8 mM glucose for 30 min. The supernatant fractions were collected to measure the secreted insulin. The insulin levels were measured with the Millipore Rat / Mouse Insulin (Cat. # EZRMI-13K), as what indicated by the manufacturer’s instructions.

### Somatostatin measurement

The somatostatin secretions were preformed similar to previously described in our group^7,9,24^. Fifty day 7 expanded islet clusters or ten primary islets for each group were starved in 1 mM glucose MKRBB buffer for 1 hour at 37°C, then treated with 20 mM glucose and 100 nM UCN3, whereas the control groups were treated with 1 mM glucose for 1 hour. The supernatant fractions were collected for somatostatin measurement using the Phoenix Pharmaceuticals ELISA kit (EK-060-03), as what indicated by the manufacturer’s instructions.

### Glucagon measurement

The glucagon secretions were preformed similar to previously described in our group^7,9,24^. Fifty day 7 expanded islet clusters or ten primary islets for each group were starved in 12 mM glucose MKRBB buffer for 1 hour at 37 °C, and then treated with 1 mM glucose and 25 mM Arginine, whereas the control groups were treated with 5 mM Arginine for 1 hour. The supernatant fractions were collected to measure the secreted glucagon. The glucagon levels were measured with the Cloud-Clone Corp Mouse Glucagon (Cat. CEB266Mu), as what indicated by the manufacturer’s instructions.

### Statistical analysis

All the qRT-PCR data were performed independently for at least three times. Statistical analyses were performed using GraphPad Prism 7 software. Experimental data were performed using unpaired two-tailed Student’s t-test. Results are presented as mean ± SEM; P < 0.05 was considered statistically significant.

## Acknowledgements

This work was supported by funding from the National Science Fund for Excellent Young Scholars Grant (81822008 to X.Y.), the National Natural Science Foundation of China (31671197 to X.Y., 81773704 to J.-P.S., 31970779 to H.-L.H., and 81173086 to J.A.), National Key Research and Development Program of China (2019YFA0111400 to H-.L.H.), and the Shandong Key Research and Development Program (2017GSF18138 to J.A.).

## Conflict of interest

The authors declare that they have no conflict of interest.

## Author contributions

X.Y. started the idea for in vitro expansion of islet tissues since 2010. X.Y. and J.-P.S. supervised the overall project design and execution. H.-L.H. provided key experience and protocols for in vitro 3D tissue culture and initial components advices. X.Y., J.-P.S. and H.-L.H. initiated the project. J.-P.S. organized the screening idea. X.Y., J.-P.S. and J.-Y.L. designed the screening details for chemicals and other factors. X.Y., J.-P.S., J.-Y.L. and J.C. designed the cell culture and all other experiments. J.-Y.L., J.C., Y.-Q.D., W.P., X.Y. and J.-P.S. participated in data analysis and interpretation. Z.Z., Y.-Q.D. and R.-J.C. provided the new material preparation methods. J.W., Z.Y., F.Y. and J.A. provided insightful idea. Y.-F.X., H.L., W.-T.A. and J.W. participated in molecular biology and animal experiments. X.Y. and J.-P.S. wrote the manuscript. H.-L.H., R.-J.C., J.-Y.L., J.C., Y.-Q.D. and W.P. revised manuscript and provided key insights for discussion.

## Contact for reagent and resource sharing

Further information and requests for resources and reagents should be directed to and will be fulfilled by the Lead Contact, Professor Xiao Yu, Jin-Peng Sun or Hui-Li Hu.

