## Supplementary Information for "In vitro pancreatic islet cluster expansion facilitated by hormones and chemicals"

### Supplementary Fig 1

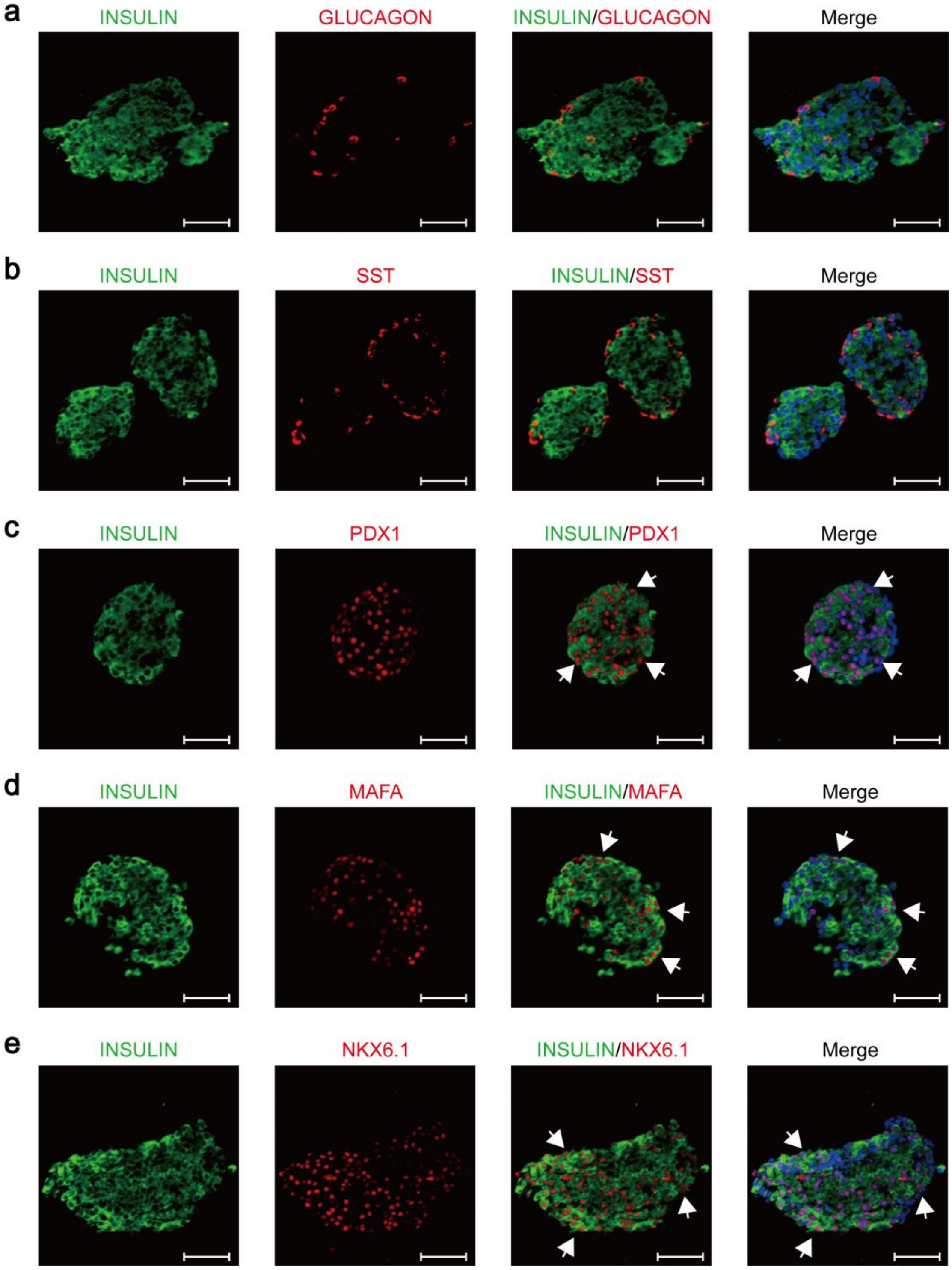

**Supplementary Fig. S1: Immunostaining for pancreatic islet cell markers of in vitro primary islets. Related to Fig. 1.**

**a** Immunofluorescence staining for INSULIN and GLUCAGON in primary islets derived from pregnant mice; the scale bar represents 50 μm.

**b** Immunofluorescence staining for INSULIN and SOMATOSTATIN in primary islets derived from pregnant mice; the scale bar represents 50 μm.

**c** Immunofluorescence staining for INSULIN and PDX1 in primary islets derived from pregnant mice; the scale bar represents 50 μm.

**d** Immunofluorescence staining for INSULIN and MAFA in primary islets derived from pregnant mice; the scale bar represents 50 μm.

**e** Immunofluorescence staining for INSULIN and NKX6.1 in primary islets derived from pregnant mice; the scale bar represents 50 μm.

### Supplementary Fig 2

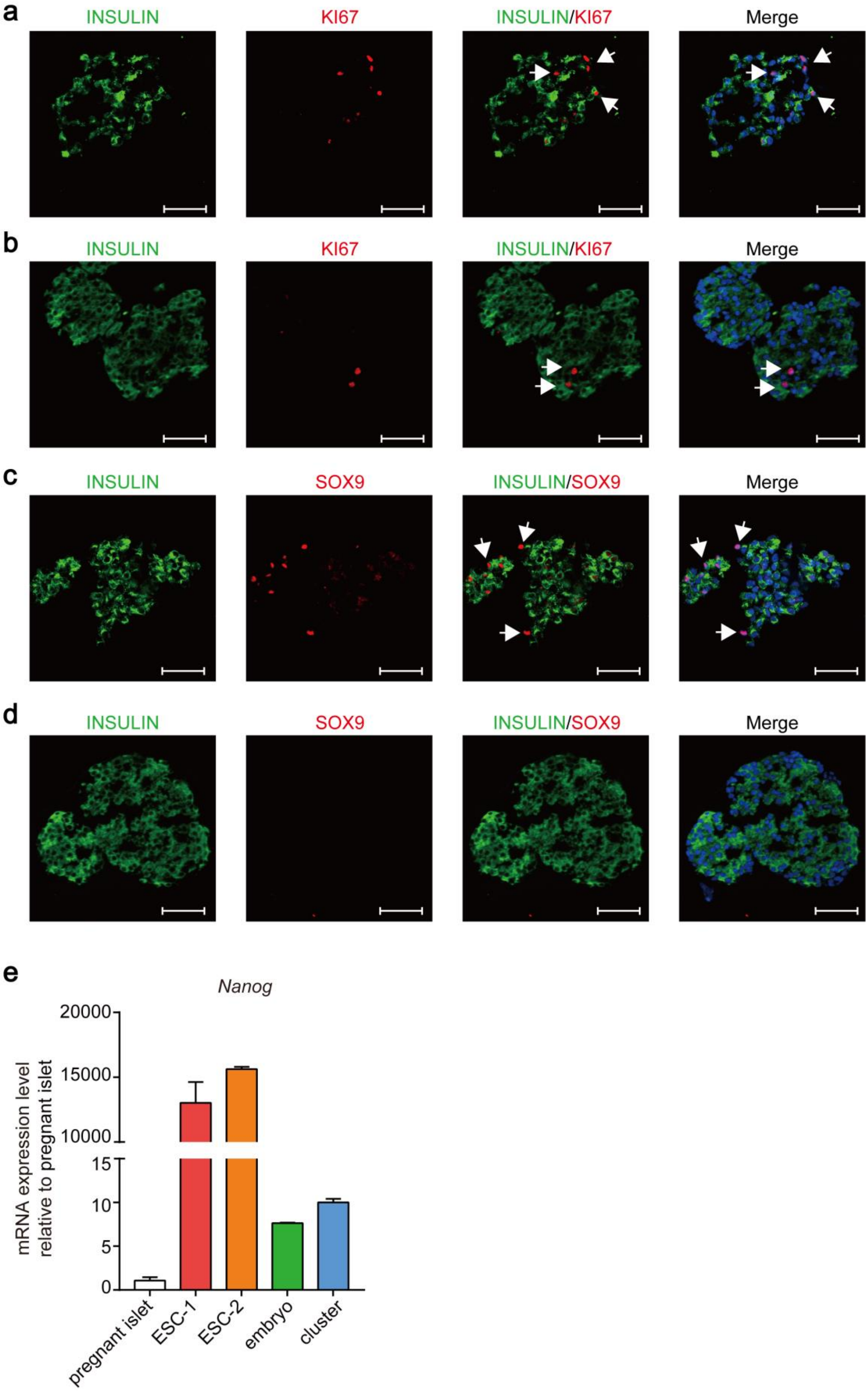

**Supplementary Fig. S2: Further analysis of gene expression in expanded clusters compared to primary islets. Related to Fig. 5.**

**a** Immunofluorescence staining for KI67 in expanded pancreatic islet clusters derived from pregnant mice; the scale bar represents 50 μm.

**b** Immunofluorescence staining for KI67 in primary islets derived from pregnant mice; the scale bar represents 50 μm.

**c** Immunofluorescence staining for SOX9 in expanded pancreatic islet clusters derived from pregnant mice; the scale bar represents 50 μm.

**d** Immunofluorescence staining for SOX9 in expanded pancreatic islet clusters derived from pregnant mice; the scale bar represents 50 μm.

**e** Comparison of the Nanog expression. qRT-PCR analysis of the Nanog expression in different cells and tissues. ESC-1 and ESC-2: two populations of embryonic stem cells of C57BL/6 mice; embryo: E11.5 embryo of C57BL/6 mice; cluster: day 7 expanded islet clusters derived from pregnant mice

### Supplementary Tables

#### Supplementary Table S1. Primer sequences used in this study

| Primers | Sequences |
|---|---|
| Gcg Forward for mouse cDNA: | GGCACATTCACCAGCGACTA |
| Gcg Reverse for mouse cDNA: | GAGAAGGAGCCATCAGCGTG |
| Ins1 Forward for mouse cDNA: | TGGACTATAAAGCTGGTGGGC |
| Ins1 Reverse for mouse cDNA: | TGTGTAGAAGAAGCCACGCT |
| Ins2 Forward for mouse cDNA: | GCTATCCTCAACCCAGCCTAT |
| Ins2 Reverse for mouse cDNA: | CTCCAGTGCCAAGGTCTGAA |
| Mafa Forward for mouse cDNA: | GGCACATTCTGGAGAGCGAG |
| Mafa Reverse for mouse cDNA: | TCACAGAAAGAAGTCGGGTGC |
| Mafb Forward for mouse cDNA: | TACCAGCAGATGAACCCCGA |
| Mafb Reverse for mouse cDNA: | GCTAGTGGGTAGCTGTTGCG |
| Sst Forward for mouse cDNA: | CCACCGGGAAACAGGAACTG |
| Sst Reverse for mouse cDNA: | TTGCTGGGTTCGAGTTGGC |
| Ngn3 Forward for mouse cDNA: | ACGCAATTTACTCCAGGCGA |
| Ngn3 Reverse for mouse cDNA: | GAGGCGCCATCCTAGTTCTC |
| Hlxb9 Forward for mouse cDNA: | GAACCTCTTGGGGAAGTGCC |
| Hlxb9 Reverse for mouse cDNA: | TCTTTGGCCTTTTTGCTGCG |
| Gata4 Forward for mouse cDNA: | AGATCGCGCCGGTTTTCTG |
| Gata4 Reverse for mouse cDNA: | GATCACCCACCGGCTAAAGA |
| Gata6 Forward for mouse cDNA: | TTTCCGGCAGAGCAGTAAGAG |
| Gata6 Reverse for mouse cDNA: | GAAACGCTTTGGCAGGCAC |
| Sox9 Forward for mouse cDNA: | CGGAACAGACTCACATCTCTCC |
| Sox9 Reverse for mouse cDNA: | GCTTGCACGTCGGTTTTGG |
| Ki67 Forward for mouse cDNA: | CAAGGCGAGCCTCAAGAGATA |
| Ki67 Reverse for mouse cDNA: | TGTGCTGTTCTACATGCCCTG |
| Nanog Forward for mouse cDNA: | AGGACAGGTTTCAGAAGCAGA |
| Nanog Reverse for mouse cDNA: | CCATTGCTAGTCTTCAACCACTG |

| | |
|---|---|
| Ccnb1 Forward for mouse cDNA: | GCGTGTGCCTGTGACAGTTA |
| Ccnb1 Reverse for mouse cDNA: | CCTAGCGTTTTTGCTTCCCTT |
| Ccnd1 Forward for mouse cDNA: | TGACTGCCGAGAAGTTGTGC |
| Ccnd1 Reverse for mouse cDNA: | CTCATCCGCCTCTGGCATT |
| Pcna Forward for mouse cDNA: | TTGCACGTATATGCCGAGACC |
| Pcna Reverse for mouse cDNA: | GGTGAACAGGCTCATTCATCTCT |
| Pdx1 Forward for mouse cDNA: | CCTTTCCCGAATGGAACCGA |
| Pdx1 Reverse for mouse cDNA: | GGGCCGGGAGATGTATTTGT |
| Sox17 Forward for mouse cDNA: | CCAAAGCGGAGTCTCGCAT |
| Sox17 Reverse for mouse cDNA: | GCCTAGCATCTTGCTTAGCTC |
| Cdk4 Forward for mouse cDNA: | GCCACTCGATATGAACCCGT |
| Cdk4 Reverse for mouse cDNA: | CACAGACATCCATCAGCCGT |
| Gcg Forward for rat cDNA: | TTCACAGGGCACATTCACCA |
| Gcg Reverse for rat cDNA: | CTATGGCGACTTCTTCCGGG |
| Ins1 Forward for rat cDNA: | ACCCTAAGTGACCAGCTACAATC |
| Ins1 Reverse for rat cDNA: | CGGGTCCTCCACTTCACGAC |
| Ins2 Forward for rat cDNA: | ACCTTTGTGGTTCTCACTTGGT |
| Ins2 Reverse for rat cDNA: | CAGTGCCAAGGTCTGAAGGTCA |
| Mafa Forward for rat cDNA: | GACAAGTTTGCGCAGGCCG |
| Mafa Reverse for rat cDNA: | GTATTCACCGTTCTCGGGGC |
| Mafb Forward for rat cDNA: | GCAACGGTAGTGTGGAGGAC |
| Mafb Reverse for rat cDNA: | GAGCTGCGTCTTCTCGTTCT |
| Sst Forward for rat cDNA: | GCTACTGGAGTCGTCTCTGC |
| Sst Reverse for rat cDNA: | GGCATCGTTCTCTGTCTGGT |
| Ngn3 Forward for rat cDNA: | GTTCCAATTCCACCCCACCT |
| Ngn3 Reverse for rat cDNA: | CGCAGGGTCTCGATCTTTGT |
| Hlxb9 Forward for rat cDNA: | CGGCGCTTTCCTACTCGTAT |
| Hlxb9 Reverse for rat cDNA: | TCCCCAAGAGGTTCGATTGC |
| Gata4 Forward for rat cDNA: | TGAATGGTATCAACCGGCCC |

| | |
|---|---|
| Gata4 Reverse for rat cDNA: | TTTGAATCCCCTCCTTCCGC |
| Gata6 Forward for rat cDNA: | TCATCACCACCCGACCTACT |
| Gata6 Reverse for rat cDNA: | GCATGCGTTGCACAGGTAAT |
| Sox9 Forward for rat cDNA: | CACAAGAAAGACCACCCCGA |
| Sox9 Reverse for rat cDNA: | TGCACGTCTGTTTTGGGAGT |
| Ki67 Forward for rat cDNA: | ACAGGGCTTAGGAAACAGTCC |
| Ki67 Reverse for rat cDNA: | GGTTCTAACTGGTCTTCCTGGT |
| Nanog Forward for rat cDNA: | AAGTCCCTTCCCTTGCCGT |
| Nanog Reverse for rat cDNA: | CTCGGGACCAGACAGCTTTAG |
| Ccnb1 Forward for rat cDNA: | GGGTGTCTTCTCAGATCGGC |
| Ccnb1 Reverse for rat cDNA: | TCCACAGGTTTTGGTAGGGC |
| Ccnd1 Forward for rat cDNA: | TCAAGTGTGACCCGGACTG |
| Ccnd1 Reverse for rat cDNA: | CTACTTGGTGACTCCCGCCT |
| Pcna Forward for rat cDNA: | GACGGGGTGAAGTTTTCTGC |
| Pcna Reverse for rat cDNA: | GACAGTGGAGTGGCTTTTGTG |

**Supplementary Table S2. Antibodies used in this study**

| **Antibodies** | **Source** | **Identifier** |
|---|---|---|
| Anti-INSULIN | Santa | sc-9168 |
| Anti-GLUCAGON | Sigma | G2654 |
| Anti-SOMATOSTATIN | Abcom | ab30788 |
| Anti-PDX1 | Abcom | ab47267 |
| Anti-SOX9 | Abcom | ab185966 |
| Anti-NKX6.1 | Abcom | ab221549 |
| Anti-MAFA | Abcom | Ab26405 |
| Anti-KI67 | Cell Signaling Technology | D3B5 |
| Donkey anti-Mouse IgG (H+L) ( secondary antibody) | Invitrogen | A10037 |
| Donkey anti- Rabbit IgG (H+L) ( secondary antibody) | Invitrogen | A21206 |
| Donkey anti-Rat IgG(H + L) (secondary antibody) | Abcam | ab150154 |

**Supplementary Table S3. Chemicals, Peptides, and Recombinant Proteins used in this study**

| Reagents | Source | Identifier |
|---|---|---|
| TRIzol | Thermo | 15596-026 |
| Cell Recovery Solution | Corning | 354253 |
| Matrigel | Corning | 356231 |
| GlutaMax | Thermo | 35050-061 |
| Penicillin-Streptomycin | Biological Industries | 03-031-1B |
| recombinant human EGF | Peprotech | AF-100-15 |
| recombinant human FGF10 | Peprotech | 100-26 |
| CHIR-99021 | BioGems | 2520691-1MG |
| N-Acetylcysteine | Sigma-Aldrich | A9165-25G |
| Nicotinamide | Sigma-Aldrich | N0636 |
| B27 Supplement (minus Vitamin A) | Thermo | 12587-010 |
| gastrin-1, human | MedChemExpress | HY-P1097 |
| A83-01 | Adooq Bioscience | A12358 |
| Y-27632 | Adooq Bioscience | A11001 |
| Forskolin | TargetMol | T2939 |
| Exendin4 | ChinaPeptides | |
| 5-Iodotubercidin | Adooq Bioscience | A13948 |
| L-Ascorbic acid | Sigma-Aldrich | A4544 |
| Triton X-100 | Solarbio | T8200 |
| Donkey Serum | Solarbio | SL050 |
| DAPI | Beyotime | C1002 |
